# A conserved ATG2-GABARAP interaction is critical for phagophore closure

**DOI:** 10.1101/624627

**Authors:** Mihaela Bozic, Luuk van den Bekerom, Beth A. Milne, Nicola Goodman, Lisa Roberston, Alan R. Prescott, Thomas J. Macartney, Nina Dawe, David G. McEwan

## Abstract

The intracellular trafficking pathway, macroautophagy, acts as a recycling and disposal service that can be upregulated during periods of stress, to maintain cellular homeostasis. An essential transition point in the pathway is the sealing of the immature phagophore to form an autophagosome, isolating unwanted cargo prior to lysosomal degradation. However, little mechanistic detail is known about phagophore closure. Human ATG2A and ATG2B proteins, through their interaction with WIPI proteins, are thought to be key players during phagophore closure. We have identified a highly-conserved motif driving the interaction between human ATG2 and GABARAP proteins that is in close proximity to the ATG2-WIPI4 interaction site. We show that the ATG2-GABARAP interaction mutants are unable to close phagophores resulting in blocked autophagy, similar to ATG2A/ATG2B double knock-out cells. In contrast, the ATG2-WIPI4 interaction mutant fully restored phagophore closure and autophagy flux, similar to wild type ATG2. Taken together, we provide new mechanistic insights to the requirements for ATG2 function at the phagophore and suggest that an ATG2-GABARAP interaction is essential for phagophore closure, whereas ATG2-WIPI4 interaction is dispensable.

## Introduction

How our cells ability to deal with a wide variety of cellular stresses is depends on two quality control pathways – namely the ubiquitin proteasome and the autophagosome to lysosome (macroautophagy) pathways. Both act in concert to ensure that our cells maintain their homeostasis. Macroautophagy (henceforth autophagy) is a multi-step process that requires the initiation and formation of a phagophore that grows and surrounds cargo to be degraded. The phagophore eventually seals to form a double-membraned vesicle, termed autophagosome. The autophagosome is then transported to, and fuses with, the lysosome where the inner autophagosomal membrane along with the cargo contents are degraded and recycled back to the cell (reviewed in [1]). This provides an intracellular pool of amino acids and lipids that the cell can utilize under periods of stress. Autophagy inducing stresses include amino acid/growth factor starvation (non-selective, bulk autophagy), mitochondrial depolarization[2, 3], pathogen invasion [4] and protein aggregate accumulation[5] (selective autophagy). In all cases, the inclusion of the cargo within the growing phagophore, and eventually the autophagosome, serves to isolate potentially cytotoxic material from the surrounding intracellular environment.

The molecular machinery involved in autophagosome formation is extensive and, for the most part, highly conserved. More than 30 ATG (<u>**A**</u>u<u>**T**</u>opha<u>**G**</u>y) genes regulate all stages of autophagosome formation; from initiation, cargo selection, transport to fusion with the lysosome. In higher eukaryotes, several kinase complexes, as well as ubiquitin-like conjugation machinery, are required for the initiation and expansion of the autophagosome. For example, the initiation kinase complex consists of ULK1/ATG13/ATG101/FIP200 and the lipid kinase complex VPS34/Beclin1/ATG14L1/p150 [6–8]. To grow the autophagosome and to recruit cargo requires the ubiquitin-like conjugation machinery, consisting of ATG7 (E1-like) ATG3 and ATG10 (E2-like) and ATG12-ATG5-ATG16L1 (E3-like complex), which are responsible for the conjugation of ubiquitin-like MAP1LC3/GABARAPs (mammalian homologues of yeast Atg8) to phosphatidylethanolamine (PE) on the growing phagophore membrane [9]. LC3/GABARAP proteins, once conjugated to PE, can localize to both the inner and outer autophagosomal membrane. This allows the ATG8s to interact with proteins containing an <u>**L**</u>C3 <u>**i**</u>nteraction <u>**r**</u>egion/<u>**G**</u>ABARAP <u>**I**</u>nteraction <u>**M**</u>otif (**LIR/GIM**), linking the phagophore to cargo or the phagophore/autophagosome to the cellular transport and fusion machinery [10–13]. The majority of LIR motifs contain a core Θ-X_1_-X_2_-Γ motif, where Θ is an aromatic residue (W/F/Y), and Γ is a large hydrophobic residue (L/V/I). In addition, acidic and/or phosphorylatable serine/threonine residues N- and C-terminal of the core LIR sequence can contribute to the stabilization of LIR-ATG8 interactions [14–16].

Despite a surge in our understanding of the mechanisms involved in autophagy, there are still questions pertaining as to how the double-membrane phagophore closes and seals to form the autophagosome. In particular, the molecular components and how they interact are relatively unknown. For example, in yeast, Vps21 (Rab5-related GTPase) and Rab5 influence phagophore closure [17, 18]. The mammalian ATG8 protein GATE-16 (GABARAP-L2) has been shown to be involved in the later stages of autophagosome biogenesis [19] and its N-terminal extension can promote membrane fusion events, hinting at a possible role during phagophore closure[20]. However, a recent study where LC3 and GABARAPs were knocked-out, indicated that LC3/GABARAPs were not required for phagophore closure [21]. A mutant form of ATG4B (C74A), the cysteine protease responsible for LC3 and GABARAP priming and removal from the autophagosomal membrane, prevents LC3 and GABARAP lipidation and results in an increased number of unsealed phagophore membranes [22]. In addition to core autophagy proteins, a component of the ESCRT-III endocytic machinery, CHMP2A, regulates the separation of inner and outer phagophore membranes [23].

One intriguing example of the role of ATG proteins during phagophore closure is the poorly understood ATG2 proteins, ATG2A and ATG2B. Mammalian ATG2s are >1900 amino acids in length and share approximately 40% amino acid sequence homology but are only 13% similar to the single isoform of *S.cerevisiae* Atg2 and 24-26% to the *D.melanogaster* Atg2, indicating a potential divergence of function. Indeed, the reconstitution of human ATG2A in yeast Δ*atg2* cells is not sufficient to restore the autophagy defects [24]. In yeast, Atg2 constitutively interacts with Atg18 at phosphatidylinositol-3-phosphate (PtIns3P)-rich membrane regions and tethers pre-autophagosomal membranes to the endoplasmic reticulum for autophagosome formation [25, 26]. Mammalian homologues of yeast Atg18 are the WIPI proteins (WIPI1-4) that are involved in various stages of autophagosome formation [27–29]. ATG2A and ATG2B preferentially interact with WIPI4 (WDR45) through a conserved Y/HFS motif [29–31]. Simultaneous depletion of both ATG2A and ATG2B results in the accumulation of small, open immature phagophore structures [32, 33]. The depletion of WIPI4 also causes open phagophore structures but they are morphologically dissimilar to those generated after ATG2A/B depletion [29]. Interestingly, previous studies have not, despite mapping the ATG2-WIPI4 interaction, shown whether this interaction is required for the restoration of autophagy flux in ATG2A/B depleted cells [29–31]. Herein, CRISPR/Cas9 was used to generate GFP-ATG2A knock-in cells as a tool to address the endogenous localization and interaction of human ATG2A. We have identified a direct interaction between the GABARAP family of mammalian ATG8 proteins and ATG2A and ATG2B that is mediated through a highly conserved LIR/GIM sequence. Surprisingly, the newly identified LIR/GIM sequence in ATG2A and ATG2B is approximately 30 amino acids N-terminal of the WIPI4 interaction motif and represent independent interaction sites in the C-terminus of human ATG2s. Using reconstituted ATG2A/2B double knockout cells, we show that the disruption of ATG2-WIPI4 interaction had no discernible effects on phagophore closure and autophagy flux, whereas mutation of the LC3/GABARAP interaction motif on ATG2 completely blocked phagophore closure and autophagy flux. Taken together, these data provide new insights into essential ATG2 interactions during autophagosome biogenesis.

## Results and Discussion

### Endogenous GFP-tagged ATG2A co-localizes and co-precipitates with GABARAP

In order to study the function of endogenous ATG2 proteins, we generated GFP-tagged ATG2A knock-in U2OS cells using CRISPR/Cas9 (Figure EV1A-1C). Under complete, nutrient-rich conditions (**CM**), GFP-ATG2A showed a dispersed localization, with little overlap with LC3B (Figure 1A, **Upper panels**). However, upon starvation we observed ring and punctate structures form that localized in close proximity to LC3B positive vesicles (Figure 1A, **Lower panels**). Endogenous ATG2B co-localized with GFP-ATG2A on both the punctate and ring-like structures observed (Figure 1A, **lower panels**). Endogenous GFP-ATG2A co-localized with early autophagy marker proteins WIPI2 (Figure 1B) and ATG16L1 (Figure 1C) at LC3B positive structures formed under starvation conditions. In addition, GABARAP-L1 was also present on GFP-ATG2A/LC3B positive structures under starvation conditions (Figure 1D). Given the presence of both GABARAP-L1 and LC3B co-localizing with GFP-ATG2A, we were curious as to whether we could co-precipitate either or both using GFP-ATG2A as bait. Using U2OS WT (control) or GFP-ATG2A U2OS cell under CM or starvation conditions, we immunoprecipitated GFP-ATG2A. WIPI4, a cognate ATG2 interaction partner [29–31], co-precipitated with GFP-ATG2A under both CM and starvation conditions (Figure 1E). We could not detect endogenous LC3B in GFP-ATG2A immunoprecipitates; but we detected increased co-precipitation of GABARAP, under starvation conditions (Figure 1E). Indeed, both ATG2A (Figure EV1D) and ATG2B (Figure EV1E) were able to co-precipitate with GFP-tagged GABARAP but not with GFP-LC3B when overexpressed in HEK293T cells. Taken together, this indicates that endogenously tagged ATG2A is active in the autophagy pathway and both ATG2A and ATG2B may functionally interact with the GABARAP family of mammalian ATG8 proteins.

**Figure 1.**
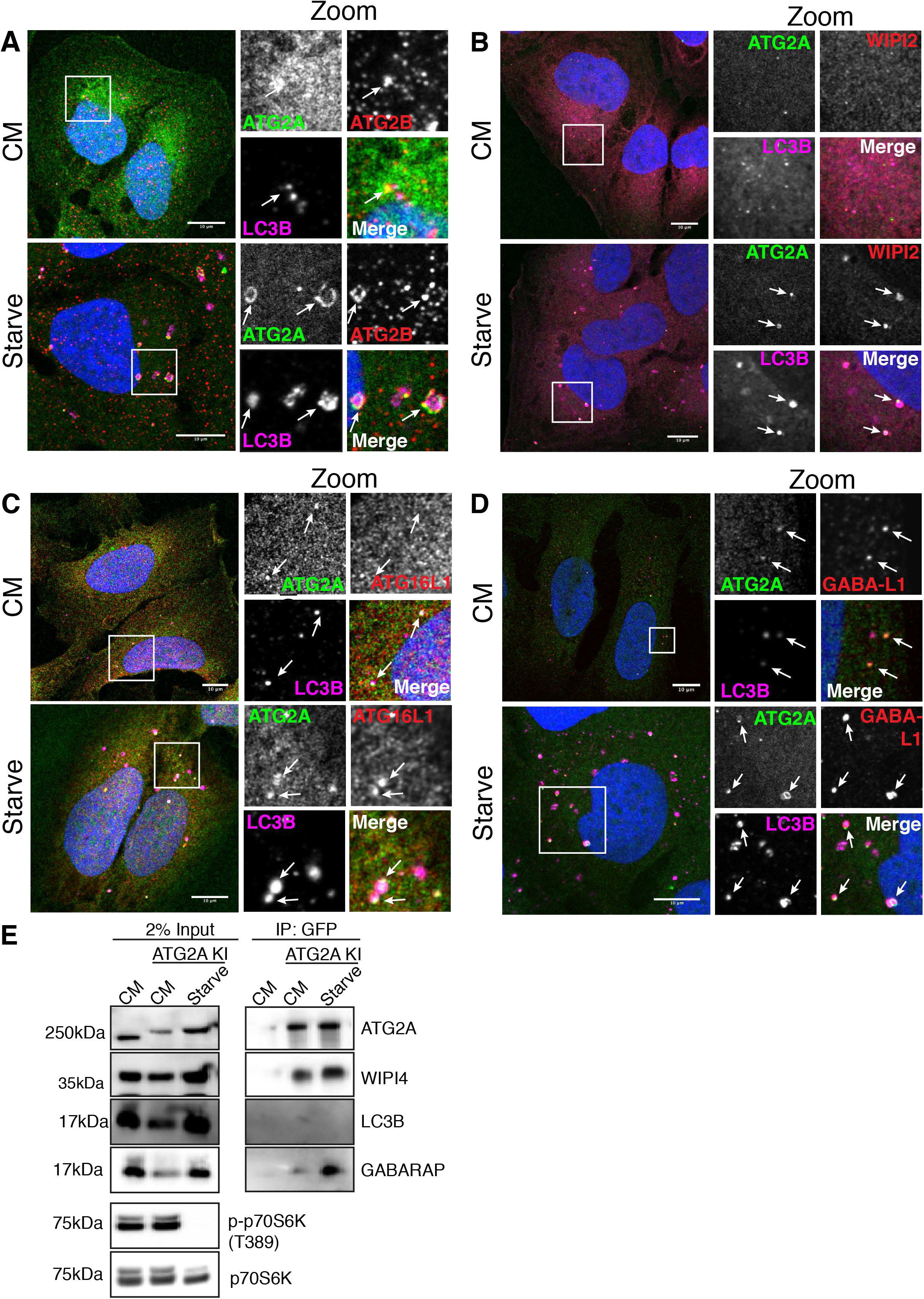
CRISPR/Cas9 GFP-tagged ATG2A localizes to early autophagy membranes. **(A)** U2OS cells modified to expressed endogenous GFP-tagged ATG2A (green) were grown in complete media (CM) or starved (EBSS) for 2 h before fixation and immunostaining with antibodies against ATG2B (red) and LC3B (magenta) and analysed by confocal microscopy. Arrows mark ATG2A/LC3B/ATG2B positive structures. Scale bar 10µm. **(B)** Cells were treated as in **(A)** and stained with anti-WIPI2 (red) or **(C)** anti-ATG16L1 (red) or **(D)** GABARAP-L1 (red). Arrows mark structures of interest. Scale bar 10µm. **(E)** U2OS WT or U2OS GFP-ATG2A knock-in (KI) cells were grown in CM or starvation media for 2 h, lysed and incubated with anti-GFP nanobody beads coupled to agarose to immunoprecipitate (IP) GFP-ATG2A. IP samples and 2% input lysates were run on 4-12% gradient gel and processed for western blotting. Anti-ATG2A, anti-WIPI4 and anti-LC3B, anti-GABARAP (pan) were used to probe for the presence/absence of autophagy proteins in the immunoprecipitated samples. p-p70S6K (T389) was used as a marker for starved cells and total p70S6K as loading control.

### Identification of a conserved LC3 interaction region sequence in ATG2A and ATG2B

Direct interaction with Atg8/LC3/GABARAP proteins is mediated through the presence of a LIR (<u>**L**</u>C3 <u>**I**</u>nteraction <u>**R**</u>egion) on the target protein [5, 13, 34–37]. ATG2 proteins have previously been shown to be part of the mammalian LC3/GABARAP interactome [38] but no direct link interaction, or consequences, has been shown. We therefore performed an *in-silico* analysis of both ATG2A and ATG2B proteins using the iLIR tool [39], as well as manual annotation, to identify potential LIRs that conform to the [E/D/S/T]-W/F/Y-X_1_-X_2_-L/I/V consensus sequence. We excluded potential LIR sequences present on secondary structures or within domains, as LIR sequences are most frequently found within disordered regions between domains [40]. We found that ATG2A contained five, and ATG2B contained six, potential LIRs (Table 1 and Figure EV2A). We then mutated the core sequence of all the potential LIR sequences in both ATG2A and ATG2B to alanine residues (Table 1) and tested the interaction using purified GST-tagged ATG8 proteins [5]. Out of the five potential LIRs present within ATG2A (Figure EV2B-E and Figure 2A **upper**) and six potential LIRs of ATG2B (Figure EV2F-G and Figure 2A **lower)**, only a single, highly conserved functional LIR was present in both ATG2A (**LIR#5**) and ATG2B (**LIR#6**). Mutation of ATG2A-LIR#5 (amino acids 1362-1365; SDE**FCIL**; Figure 2A, **mLIR**) and ATG2B-LIR#6 (amino acids 1491-1494; NDD**FCIL**; Figure 2A, **mLIR**) resulted in the loss of GST-ATG8 interaction, compared to WT proteins (Figure 2A). Next, we overexpressed ATG2A/ATG2B-WT or LIR mutants with GFP alone, GFP-LC3B or GFP-GABARAP and immunoprecipitated the GFP-tag. ATG2A-WT (Figure 2B) and ATG2B-WT (Figure 2C) co-precipitated mainly with GABARAP and this interaction was abolished when their core LIR sequence was mutated to alanine (**mLIR**; **FCIL/AAAA**; Figure 2B-2C). ATG2A had a second potential LIR sequence (aa926-929 FSTL/AAAA; **mLIR#2**). However, immunoprecipitation of GFP-tagged LC3B and GABARAP showed that Myc-ATG2A-mLIR#2 was still able to interact with GABARAP to the same extent as ATG2A-WT; whereas ATG2A-mLIR#5 abolished the interaction (Figure EV2H).

**Table 1.**
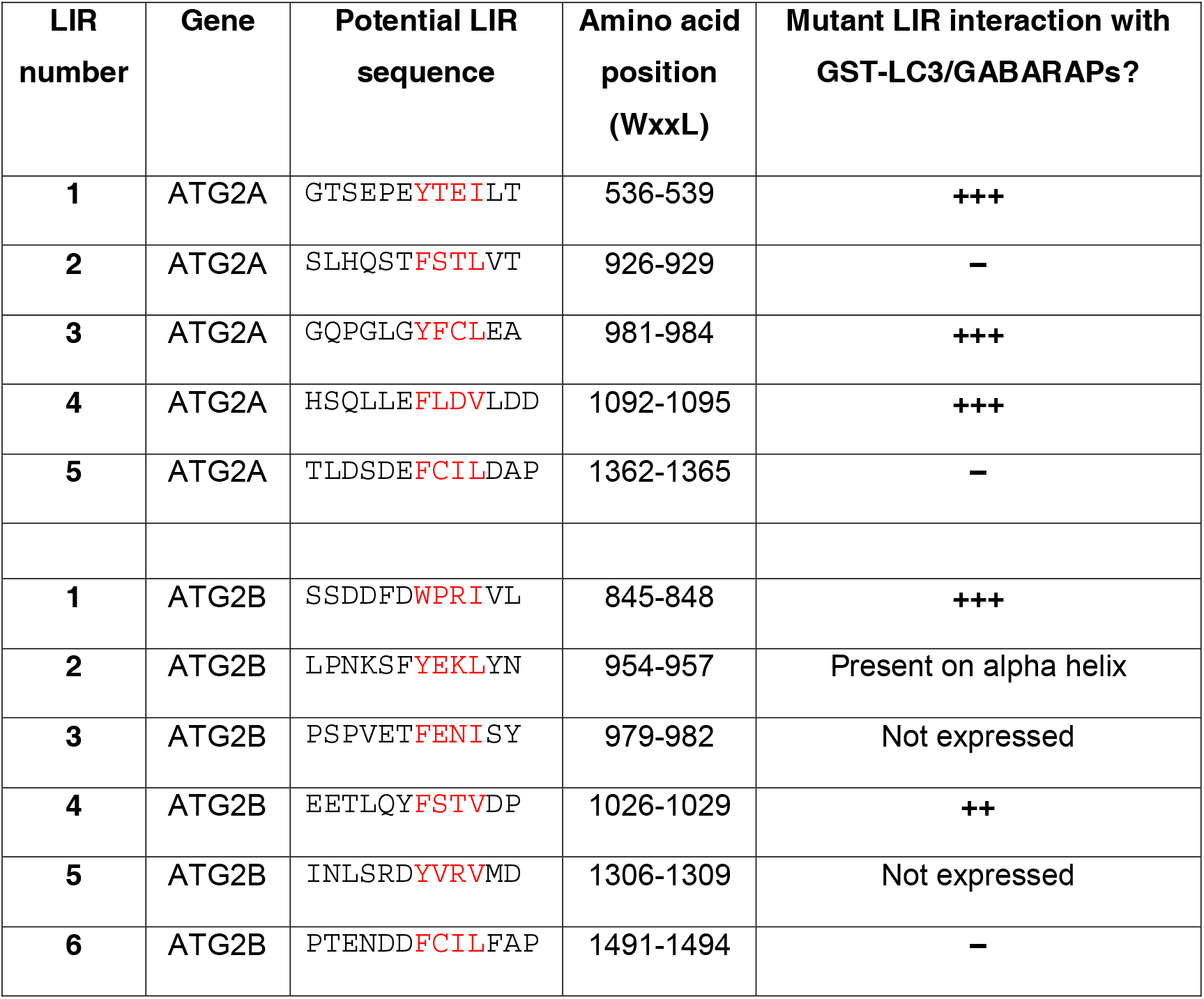
Potential LC3 interaction region sequences identified in human ATG2A and ATG2B protein sequences.

**Figure 2.**
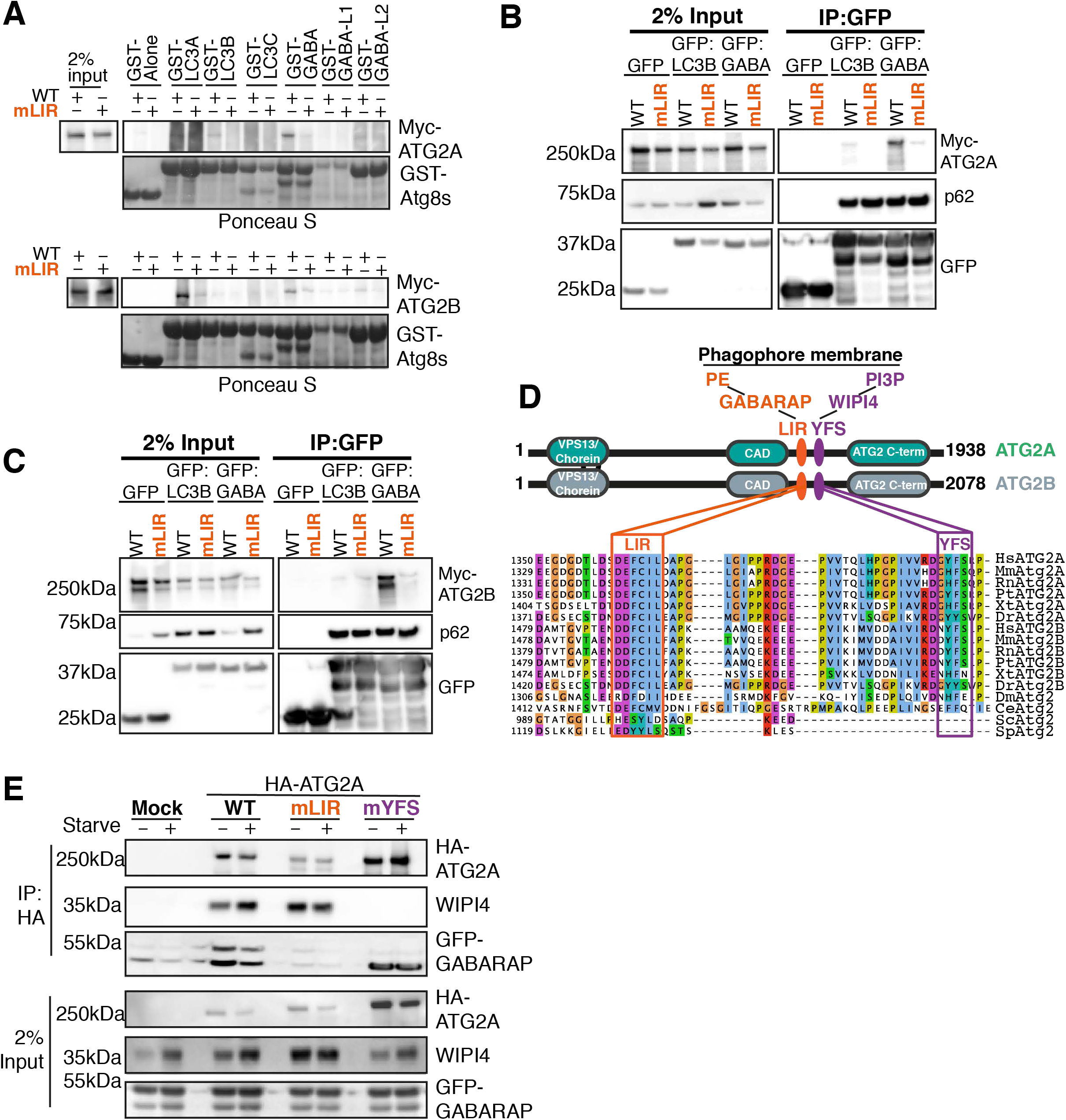
ATG2A and ATG2B contain a highly conserved LC3 Interaction Region (LIR). **(A)** Myc-tagged ATG2A-wild type (WT), ATG2A-mutant LIR (mLIR; FCIL/AAAA) (upper blots) or Myc-tagged ATG2B-WT or ATG2B-mLIR (FCIL/AAAA; lower blots) were over expressed in HEK293T cells, and lysates were incubated with purified GST alone or GST-tagged LC3A, -LC3B, LC3C, -GABARAP, -GABARAP-L1 or GABARAP-L2. Samples were spun, washed and blotted for the presence/absence of Myc tagged ATG2s using anti-Myc antibody. GST-proteins were visualised by Ponceau S staining of membranes. **(B)** Myc-tagged ATG2A-wild type (WT) and ATG2A-mLIR (orange) were co-expressed with GFP-alone, GFP-LC3B or GFP-GABARAP in HEK293T cells, lysed and anti-GFP nanobodies coupled to agarose were used to immunoprecipitate GFP-tagged proteins. Samples were subjected to western blotting and probed for the presence of Myc-ATG2A in immunoprecipitated samples. Anti-p62/SQSTM1 was used as an internal control for the immunoprecipitated samples. **(C)** As in (B) but using Myc-tagged ATG2B-WT or ATG2B-mLIR (orange) co-expressed with GFP-alone, GFP-LC3B or GFP-GABARAP. Samples were subjected to western blotting and probed for the presence of Myc-ATG2B in immunoprecipitated samples. Anti-p62/SQSTM1 was used as an internal control for the immunoprecipitated samples. **(D)** Domain structure of ATG2A (green) and ATG2B (grey) proteins. Both ATG2s contain an N-terminal VPS13/Chorein domain, ATG2 C-terminal autophagy domain (CAD motif) and ATG2 c-terminal domain. Position and sequence of the new ATG2 LIR motif that directs GABARAP interaction (orange) and the previously identified WIPI4 interaction motif (YFS; purple box). Approximately 30 amino acids separate these two motifs in human ATG2A and ATG2B. Multiple sequence alignment of multiple species using Jalview highlights the conservation of these regions. Abbreviations for species: Hs – Homo sapien; Mm – Mus musculus; Rn – Rattus norvegicus; Pt – Pan troglodytes; Xt – Xenopus tropicalis Dr – Danio rerio; Dm – Drosophila melanogaster; Ce – Caenorhabditis elegans; Sp – Schizosaccharomyces pombe; Sc – Saccharomyces cerevisiae. **(E)** HA-tagged ATG2A-WT, -mLIR (orange) and -mYFS (YFS/AAA; purple) were stably expressed in U2OS ATG2A/B double knock-out cells using retrovirus transduction. Cells were transfected with GFP-GABARAP and 24 h later grown in complete medium (CM) or starved for 2 h (EBSS). Cells were lysed and anti-HA beads were used to immunoprecipitate HA-tagged ATG2A and processed for western blot. Blots were then probed with antibodies against HA-tag (ATG2A), anti-WIPI4, and anti-GFP for the presence/absence in immunoprecipitated samples. All blots are representative of at least n=3 independent experiments.

Alignment of the amino acid sequences of ATG2 proteins from multiple species revealed that the ATG2A/2B LIR sequence is highly conserved in multiple vertebrates and invertebrates (Figure 2D, **orange box**). This includes organisms with a single ATG2 isoform; *Drosophila* melanogaster (DmAtg2), *Caenorhabditis* elegans (CeAtg2), and species with two ATG2 isoforms; *Danio* rerio (DrAtg2a/DrAtg2a) and *Xenopus* tropicalis (XtAtg2a/XtAtg2b). However, the LIR does not appear to be present in *Saccharomyces cerevisiae or Saccharomyces pombe*, indicating a potential divergence in Atg2 function (Figure 2D, **orange box**). Taken together, both ATG2A and ATG2B contain a single, highly conserved LIR motif that preferentially interacts with the GABARAP family of mammalian ATG8 proteins.

Since the first LC3 interaction region was identified in the prototypical autophagy receptor protein, p62/SQSTM1 [10], the number of functional LIR-containing proteins identified to date has grown considerably. The interaction between mammalian ATG8s and LIR containing proteins serves to control all aspects of the autophagy pathway, from cargo selection to formation, transport and fusion of the autophagosome. Not only are these interactions sequences present in mammalian, plant, fungi and invertebrate species, they are also present in a number of viral [41] and bacterial [42] proteins, potentially to aid pathogen survival and subversion of the pathway. We have identified a highly conserved LIR within both ATG2A and ATG2B that differ only in a few amino acids both N-and C-terminal of the core LIR sequence (FCIL; Figure 2D). This raises an ongoing question as to how specificity within the system is achieved, particularly in mammalian systems that are complicated by the expression of six LC3/GABARAP isoforms. We, and others [13, 43, 44], have attempted to decipher the code that dictates whether a protein with a particular LIR sequence will preferentially interact with LC3 over GABARAP. Interestingly, ATG2A and ATG2B do not conform to the recently identified GABARAP interaction motif consensus sequence (W/F-I/V-X-I/V) [13] despite preferring GABARAP over LC3 in co-immunoprecipitation from cells (Figure 1E; Figure 2B-C). Surprisingly, both ATG2A and ATG2B can also interact with LC3A (Figure 2A), however the functional consequences and the *in vivo* preference appears directed towards GABARAP proteins. The high degree of conservation of the ATG2A/B LIR sequence throughout vertebrates and invertebrates (Figure 2D) potentially indicates a conserved function, even in species with only a single ATG2 isoform, such as D.melanogaster and C.elegans. Therefore, understanding the role of ATG2-LC3/GABARAP interaction during autophagy will provide insights into ATG2s mechanism of function in multiple species.

### ATG2-WIPI interaction is not affected by mutation of the ATG2 LIR

Intriguingly, the recently described WIPI4 interaction region that contains an essential Y/HFS motif [30, 31] is approximately 30 amino acids C-terminal of the newly identified LIR sequence in both ATG2A and ATG2B (Figure 2D, **purple box**). Therefore, we wanted to test whether the interaction between ATG2, GABARAP and WIPI4 were co-dependent, or whether they represented independent interactions. The YFS motif found in ATG2A (amino acids 1395-1397; **YFS/AAA**; **mYFS**) was mutated and HA-tagged ATG2A-WT, -mLIR or - mYFS variants were immunoprecipitated from cell lysates and probed for the presence of GFP-tagged GABARAP and endogenous WIPI4. ATG2A-WT co-precipitated with both WIPI4 and GFP-GABARAP under complete media (CM) and starvation conditions (Figure 2E). ATG2A LIR mutant failed to co-precipitate GFP-GABARAP, whereas the WIPI4 interaction was unaffected (Figure 2E; **mLIR**). Conversely, mutation of the ATG2A YFS motif resulted in the loss of WIPI4 interaction but not GABARAP (Figure 2E; **mYFS**). Interestingly, the loss of the WIPI4 interaction site on ATG2A resulted in increased co-precipitation of lipidated GABARAP with ATG2A-mYFS (Figure 2E). Thus, within a 30 amino acid stretch mammalian ATG2 proteins are two distinct and independent interaction motifs that can potentially regulate ATG2 function at the growing phagophore membrane. It remains to be seen whether either or both, interactions are necessary for ATG2 function during autophagy.

### ATG2 LIR mutant but not WIPI4 mutant is sufficient to block autophagy

ATG2A can simultaneously interact with both GABARAP and WIPI4 (Figure 2E). Interestingly, previous reports that identified the WIPI4 interaction region on ATG2 [30, 31] and the ATG2 interaction site of WIPI4 [29] did not address the role of these interaction mutants during autophagy flux. Considering the close proximity of both GABARAP and WIPI4 motifs on ATG2, we wanted to dissect the individual roles of ATG2-GABARAP and ATG2-WIPI4 interactions during autophagy. Firstly, ATG2A/ATG2B double-knockout (DKO) U2OS cells were generated, that could be reconstituted with WT and mutant genes. In ATG2A/2B DKO cells, endogenous ATG2A and ATG2B were depleted compared to WT and p62/SQSTM1, lipidated mammalian ATG8s (LC3A, LC3B, LC3C, GABARAP-L1 and GABARAP-L2) were markedly increased compared to WT U2OS cells (Figure EV2I). Next, stable expression of the tandem tagged-LC3B autophagy reporter (mCherry-GFP-LC3B; [45]) in ATG2A/B DKO cells was used to assess LC3B transition from autophagosomes (GFP+ve/mCherry+ve) to autolysosomes (GFP–ve/mCherry+ve) due to GFP quenching at low pH [45]. Using confocal imaging and flow cytometry to quantify, tandem-tagged LC3B puncta in ATG2A/2B DKO cells under complete medium (CM) or starvation conditions, were both GFP and mCherry positive (Figure 3A and **quantified in 3B**). This indicated that the ATG2A/B DKO U2OS cells had impaired autophagy flux, consistent with previous work [46]. Stable reconstitution of tandem-LC3B expressing ATG2A/B DKO cells with ATG2A-WT resulted in more mCherry only positive cells/puncta in complete medium (CM) conditions that were increased upon starvation (Figure 3A and **quantified in 3B**). However, this effect was nullified using BafliomycinA1 (to prevent lysosome acidification and quenching of GFP signal) (Figure 3A and **quantified in 3B**). Surprisingly, expression of ATG2A-mLIR resulted in a complete lack mCherry only puncta/cells, indicative of impaired autolysosome formation and closely resembled the ATG2A/B DKO cells (Figure 3A and **quantified in 3B**). Unexpectedly, the ATG2A-mYFS (WIPI4 mutant) was able to fully restore autophagy flux, as indicated by increased mCherry positive puncta/cells and was comparable to the ATG2A-WT results (Figure 3A and **quantified in 3B**).

**Figure 3.**
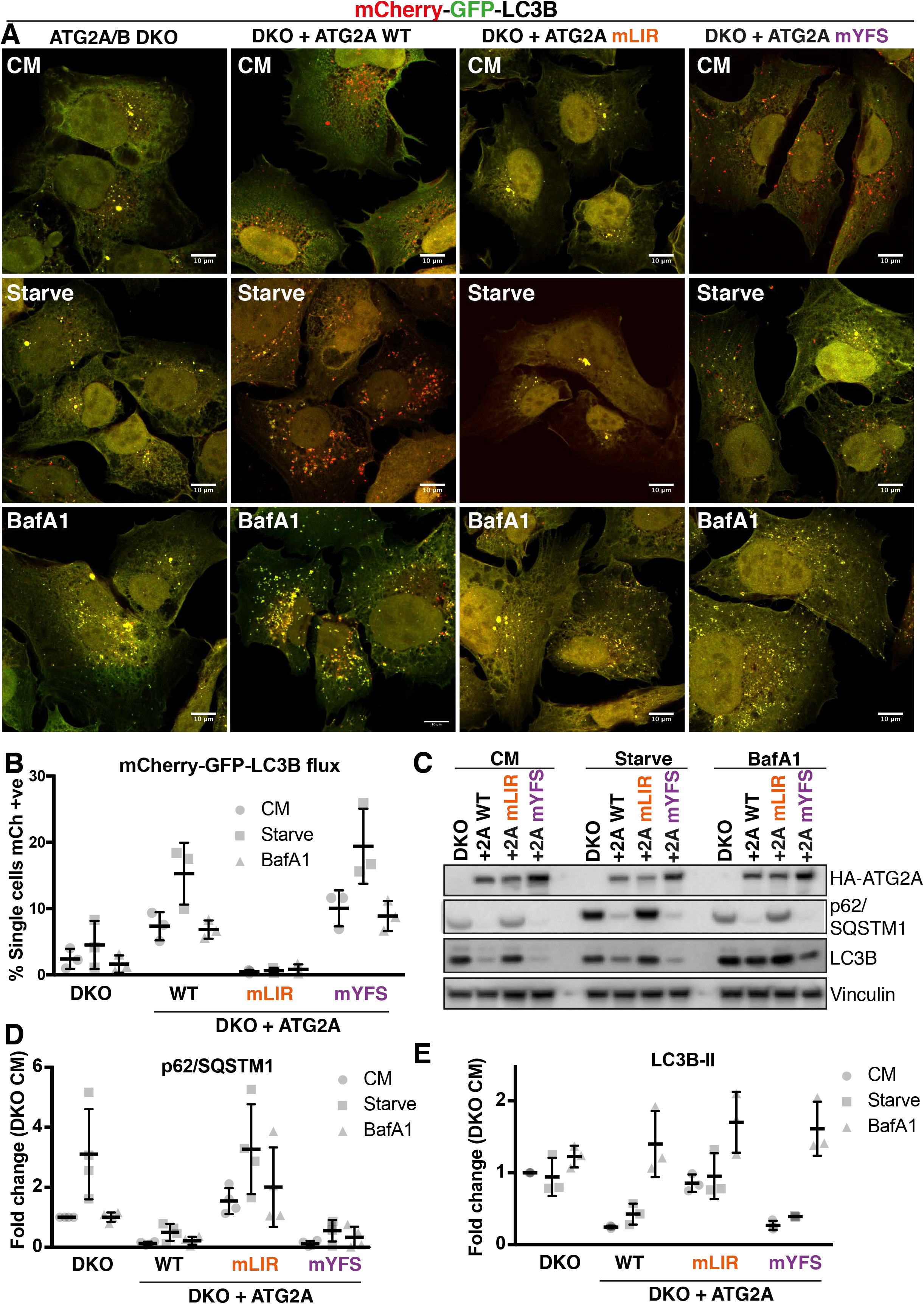
ATG2A LIR domain is essential for autophagy flux. **(A)** U2OS ATG2A/B double knock-out (DKO) CRISPR/Cas9 cells stably expressing Tandem tagged LC3B (mCherry-GFP-LC3B) were retrovirally transduced to express vector, or HA-tagged ATG2A-WT, -mLIR (FCIL/AAAA) or -mYFS (YFS/AAA). Cells were grown in complete medium (CM) or starved for 2 h (EBSS) or treated with CM plus BafilomycinA1 (200nM, 4 h), fixed and analysed by confocal microscopy. Merged images of GFP (green) and mCherry (red) channels show the presence of autophagosomes/phagophores (GFP and mCherry positive, yellow puncta) or autolysosomes (mCherry only, red puncta) Scale bar 10µm. Images are representative of n=3 independent experiments. **(B)** Quantification of (A) using flow cytometry of measuring GFP and mCherry fluorescence. Cells were gated based on GFP and mCherry fluorescence and % mCherry positive cells gated and used as an indication of autolysosome formation due to GFP quenching. Each symbol represents an n=1 independent experiment with 10,000 cells analysed per condition. Horizontal bar indicates mean ± SD. **(C)** U2OS ATG2A/B DKO cells reconstituted with vector only, HA-ATG2A-WT, -mLIR and - mYFS were stimulated with complete medium, (CM), 2 h starvation (EBSS) or 4 h BafilomycinA1 (BafA1, 200nM), lysed in total cell lysis buffer and subjected to western blot analysis. Blots were probed for the presence of HA-tag (ATG2A), p62/SQSTM1, LC3B and vinculin (loading control). **(D)** p62/SQSTM1 and LC3B-II **(E)** levels were normalized to loading control and quantified as fold change of DKO proteins levels. Each symbol represents an independent experiment. Quantification of at least n=3 independent experiments is shown. Horizontal bar represents mean ± SD.

Next, we analysed the effect of ATG2A-WT, -mLIR and -mYFS expression on both p62/SQSTM1 and LC3B protein levels, as these are autophagy substrates and are good indicators of flux [47]. Stable expression of HA-tagged ATG2A-WT in ATG2A/2B DKO cells resulted in decreased p62/SQSTM1 and LC3B-II levels, compared to DKO alone, indicating rescue of the pathway and restoration of autophagy flux (Figure 3C-E, **WT lane**). Consistent with the tandem-tagged LC3B reporter assay (Figure 3A-B), expression of ATG2A-mYFS was able to fully restore autophagy flux under nutrient-rich (CM, complete medium), starvation and bafilomycin treatment (Figure 3C-E). Excitingly, the expression of ATG2A-mLIR failed to rescue the defect in p62/SQSTM1 and LC3B-II (Figure 3C-E). In DKO plus ATG2A-WT and DKO plus ATG2A-mYFS expressing cells, LC3B was present within LAMP2 positive vesicles (lysosomes) after starvation plus BafliomycinA1 treatment, to induce autophagosome formation but halt their degradation (Figure EV3A, **open arrows**). In stark contrast, LC3B was observed juxtaposed to LAMP2 vesicles in both ATG2A/2B DKO and DKO plus ATG2A-mLIR (Figure EV3A, **closed arrows**), indicating an autophagosome maturation defect and consistent with the tandem-LC3B reporter assay (Figure 3A-B). One aspect of mammalian ATG2 function is the regulation of the size and distribution of lipid droplets (LDs) [33]. ATG2A localizes to the limiting membrane of LDs [33, 46]. Importantly, both HA-tagged ATG2A-mLIR and ATG2A-mYFS, as well as HA-ATG2A-WT, are able to localize to lipid droplets induced by oleate, a fatty acid supplement that induces the accumulation of neutral lipids into LDs (Figure EV3B-C). Therefore, disruption of either the ATG2A-GABARAP or ATG2A-WIPI4 interaction does not affect ATG2A localization to LDs. Taken together, our data shows that mutation of a conserved LC3/GABARAP interaction motif on ATG2A fails to restore the autophagy defect of ATG2A/ATG2B double knock out cells; whereas, the interaction with WIPI4 is dispensable for autophagy flux.

### ATG2-GABARAP interaction is essential for phagophore closure

Previous work has shown that early autophagy marker proteins accumulate on LC3B positive structures in ATG2A/B depleted cells [33]. Due to the similarities between the ATG2A/B DKO and ATG2A-mLIR phenotypes, we stained for the presence of several autophagy marker proteins under starvation conditions. ATG2A/2B DKO and DKO cells expressing ATG2A-mLIR exhibited large LC3B-positive/p62-positive structures (Figure EV4A, **closed arrows**) and accumulated ATG9A (Figure EV4B, **closed arrows**), WIPI2 (Figure EV4C, **closed arrows**) and ATG16L1 (Figure EV4D, **closed arrows**). In contrast, ATG2A/B DKO cells reconstituted with ATG2A-WT or ATG2A-mYFS resulted in vesicular LC3 and punctate p62/SQSTM1 structures (Figure EV4A, **open arrows**), juxtanuclear ATG9A localization (**Figure EV4B**, **open arrows**) and punctate WIPI2 (Figure EV4C, **open arrows**) and ATG16L1 (Figure EV4D, **open arrows**) consistent with restoration of efficient autophagosome biogenesis and autophagy flux.

In cells expressing ATG2A-mLIR, LC3B is lipidated (LC3B-ii; Figure 3C), early phagophore associated proteins are present (ATG9A and ATG16L1, Figure EV4B and D respectively) and the membranes contain PI3P (inferred by the presence of WIPI2, Figure EV4C). This indicates that the observed structures (Figure EV4 A-D) are phagophores or autophagosomes.

The mammalian ATG2 proteins have been suggested to function at the transition stage of phagophore to autophagosome maturation, the closure step [32, 33]. Therefore, in order to address the functional significance of the ATG2 interaction with GABARAP, used two assays to distinguish between phagophores and autophagosomes – a proteinase K protection assay (Figure 4A(i)) and Syntaxin17 (STX17) translocation (Figure 4A(ii)). Firstly, using the proteinase K limited proteolysis assay, which degrades proteins not protected within a membrane compartment (Figure 4A (i)), we tested whether the expression of ATG2A LIR mutant resulted in defective phagophore closure. ATG2A/B DKO cells and DKO cells reconstituted with ATG2A-WT, -mLIR or -mYFS (Figure 4B) were left in CM or starved for four hours in the presence of BafilomycinA1 (Starve+BafA1) to accumulate autophagosomes. Cells were then permeabilised using digitonin and incubated in buffer only, proteinase K or proteinase K plus Triton X-100 (to permeabilise membranes). Under CM conditions, the majority of the autophagy substrate p62 was degraded in all samples (Figure 4B, **upper panel)**. After starvation plus BafilomycinA1 treatment, DKO cells reconstituted with ATG2A-WT and ATG2A-mYFS showed a large proportion of p62 resistant to proteinase K degradation (Figure 4B, **lower panel**; 38% and 56% respectively). However, p62 in both DKO and DKO plus ATG2A-mLIR cells was sensitive to proteinase K digestion (Figure 4B, **lower panel**; 18% and 17% respectively), which is indicative of immature/unsealed autophagosomes.

**Figure 4.**
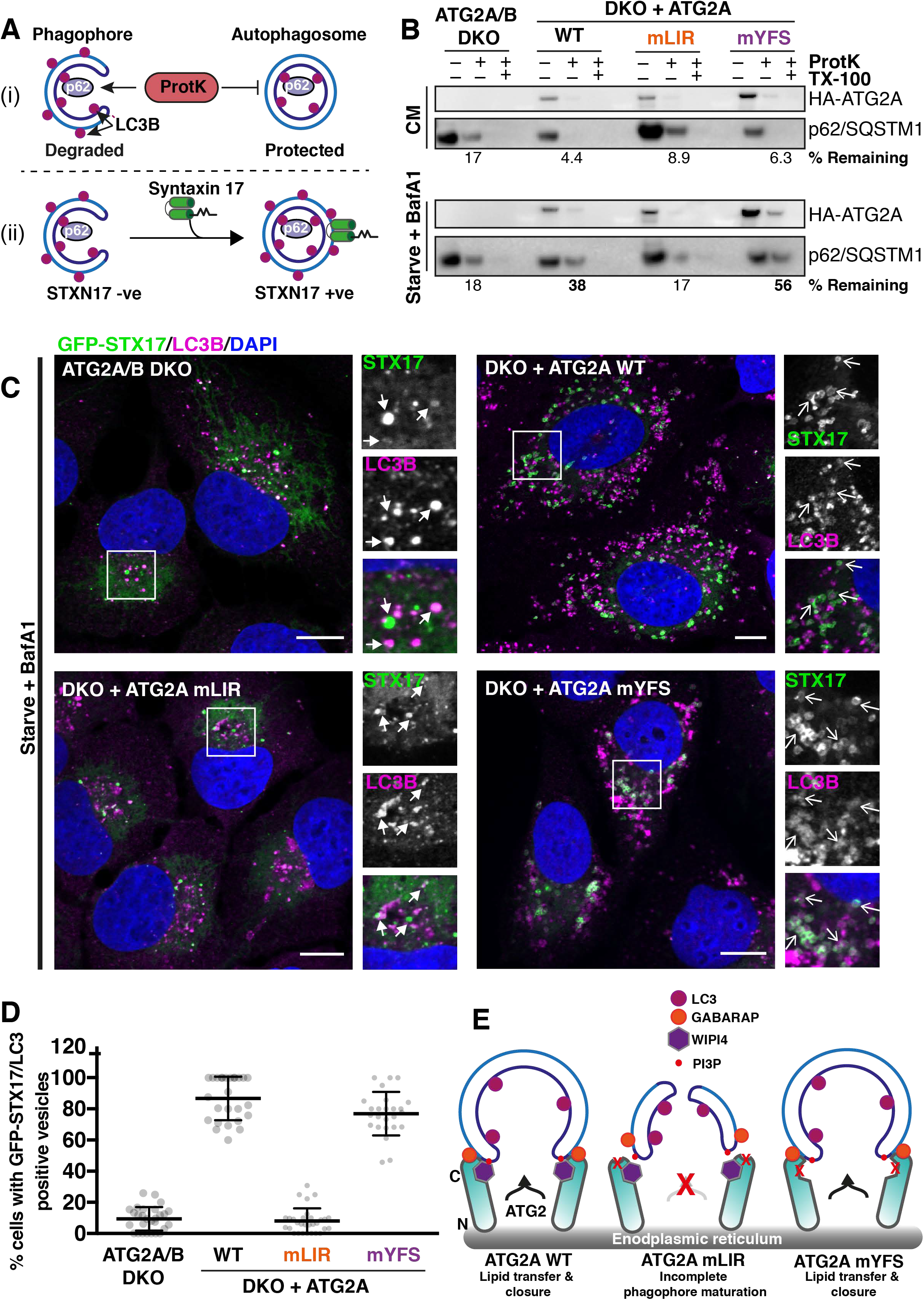
Mutation of ATG2A LIR prevents phagophore closure. **(A)** Graphical representation of proteinase K assay **(i)** showing protection of p62/SQSTM1 inside sealed autophagosomes or proteinase K sensitive p62 present within open phagophores. **(ii)** Graphical representation of sytnatxin17 (STX17) translocation to completed autophagosomes and not phagophores. Autophagosomes are identified as being both LC3B and STX17 positive vesicles. **(B)** U2OS ATG2A/B DKO cells reconstituted with vector only, HA-ATG2A-WT, -mLIR and - mYFS were stimulated with complete medium, (CM), 4 h starvation (EBSS) plus BafilomycinA1 (BafA1, 200nM) treatment. Cells were centrifuged and resuspended in PBS digitonin, spun, washed and the membrane fractions incubated with proteinase K with and without 0.1% TritonX-100. Sampels were then subjected to western blotting using anti-p62/SQSTM1 and anti-HA (ATG2A) antibodies. Percentage p62/SQSTM1 remaining was calculated using densitometry analysis. Blots are representative of n=3 independent experiments. **(C)** U2OS ATG2A/B DKO cells reconstituted with vector only, HA-ATG2A-WT, -mLIR and - mYFS and stably expressing GFP-Syntaxin17 (STX17) were stimulated starvation (EBSS) plus BafilomycinA1 (BafA1, 200nM) for 4 hours to stimulate autophagosome generation and prevent their degradation in the lysosome. Cells were fixed and immune-stained for LC3B (magenta). DAPI was included (blue) to mark the DNA/nucleus. Confocal analysis of LC3B and GFP-STX17 (green) localization was performed. Closed arrows (ATG2A/B DKO and DKO + ATG2A mLIR) highlight aggregate structures. Open arrows (DKO + ATG2A-WT and ATG2A-mYFS) highlight STX17/LC3B positive vesicles. Scale bar 10µm. **(D)** Quantification of **(C)** expressed as a percentage of cells with STX17/LC3B positive vesicles. Each symbol represents a single field of cells with 5-10 cells per field. A total of 300-600 cells were analysed over n=3 independent experiments. **(E)** Model of ATG2 function based on the current knowledge. ATG2A localizes to ER membranes and facilitates lipid transfer from the ER to the growing phagophore. ATG2 interaction with GABARAP is essential for anchoring ATG2 to growing phagophore and mutation of the GABARAP interaction region results in the formation of immature phagophores that fail to close.

Next, STX17 translocation to LC3B positive vesicles in the ATG2A-WT and mutant-expressing cells was tested. STX17 translocates from the ER to fully formed autophagosomes, but not phagophores, prior to their fusion with the lysosome (Figure 4A (ii))[48]. Stable expression of GFP-syntaxin17 in the reconstituted ATG2A/B DKO cells revealed that STX17 can efficiently localize to, and surround, LC3B positive structures in both ATG2A-WT and ATG2A-mYFS expressing cells after starvation plus bafilomycinA1 treatment (Figure 4C, open arrows and quantified in Figure 4D). Conversely, in ATG2A/B DKO and DKOs plus ATG2A-mLIR cells, GFP-STX17 localized mainly to ER and punctate structures with few GFP-STX17+ve/LC3B+ve vesicles observed (Figure 4C, closed arrows and quantified in Figure 4D). Taken together, our data suggest that a conserved GABARAP interaction motif in both mammalian ATG2A and ATG2B proteins is essential for phagophore to autophagosome transition, and surprisingly, that the WIPI4 interaction is dispensable for this function during starvation-induced autophagy.

The formation of the autophagosome and its subsequent trafficking and fusion with the lysosome is a tightly controlled pathway with a number of essential components that allows it to progress in an orderly fashion. This enables the cell to liberate amino acid and lipid stores during periods of stress, target and remove intracellular pathogens or remove cytotoxic protein aggregates from the cell. Critical to this process is the ability of the cell to form a double-membraned phagophore that grows, surrounds and isolates the material to be removed. Despite recent advances in our knowledge, the mechanisms involved in phagophore closure are poorly understood. Recent work has shown that the ESCRT-III component CHMP2A regulates the separation of inner and outer phagophore membranes [23]. In addition to the ESCRT-III machinery, TRAPPC11, a member of TRAPP complexes involved in membrane trafficking, has been shown to recruit ATG2B-WIPI4 to phagophores in an ATG9A dependent manner [49]. The depletion of TRAPC11 results in a phenotype similar to that of ATG2A/B DKO and ATG2A-mLIR [49]. The mammalian ATG2 proteins, ATG2A and ATG2B, have been shown to be essential for phagophore formation and closure [32, 33, 46] and depletion of WIPI4, a constitutive interaction partner of mammalian ATG2s, also negatively impacts on phagophore closure [29]. Herein, we have described a hitherto unidentified LC3/GABARAP interaction region on both ATG2A and ATG2B that is essential for phagophore formation and closure.

These results shed new light on the role of ATG2 during autophagosome biogenesis, and in particular, the interactions that are necessary for this process. Perhaps most surprisingly was the effect, or rather lack thereof, that the ATG2A-WIPI4 interaction mutant had on phagophore closure and autophagy flux. From yeast to fruit flies to humans, the ATG2-ATG18 (WIPI) interaction is highly conserved. In yeast, Atg2-Atg18 interaction occurs independently of Atg18 binding PI3P [26], much like the ATG2A/B interaction with WIPI4 [29–31]. Recently, yeast Atg2 has been shown to contain both N- and C-terminal membrane binding domains that help tether Atg2 to membrane contact sites [25]. Human ATG2A has several domains that determine its ability to localize to membranes. Firstly, ATG2 has an N-terminal membrane binding region that is essential for autophagosome formation [46] that has now been shown to be a lipid transport domain [50, 51]. This N-terminal lipid transport domain is thought to be essential for the transport of PE and phosphatidylserine from the ER/omegasome to the growing phagophore [50, 51]. This domain is homologous to the Vps13 lipid transport domain involved in organelle-organelle contact sites [52]. The second lipid interaction region of ATG2A is an amphipathic helix (AH; aa1750-1767), which is essential for ATG2 localization to both lipid droplets and isolation membranes and is essential for autophagy flux [46]. In addition, ATG2 has a c-terminal region (aa1830–1938 HsATG2A) that is involved in localization to lipid droplets but is dispensable for autophagy [46]. This raises an interesting question as to the role of the ATG2-WIPI4 interaction, as this was previously thought to be involved in ATG2 autophagy function. However, we have shown that ATG2-WIPI4 is dispensable for autophagosome formation and autophagy flux. Given that the ATG2A LIR mutant we identified has impaired autophagy flux (Figure 3) but can still localize to lipid droplets (Figure EV3B-C), we suggest that both the ATG2-AH and the ATG2-LIR are essential and the combination of both the LIR and AH helps to define the target membrane, allowing tethering and lipid transfer to facilitate efficient phagophore formation and autophagosome maturation (Figure 4E).

## Materials & Methods

### Antibodies

The antibodies used in this study as re as follows: Anti-GFP (Santa Cruz clone B-2, sc9996), anti-FlagM2 (SIGMA, F3165), anti-p62 (MBL, M162-3), anti-LC3B (clone 5F10 Nanotools, 0231-100/LC3-5F10) and anti-GABARAP (Abcam, ab109364) anti-ATG16L1 (MBL, PM040), anti-ATG9A (abcam, ab108338), anti-WIPI2 and anti-WIPI4 were kind gifts from Prof. Sharon Tooze, LC3A clone D50G8 (CST, #4599), anti-LC3C (D1R8V; CST # 14723), anti-GABARAP (Abcam, ab109364), anti-GATE-16 (MBL, PM038), anti-GABARAP-L1 (IF, Proteintech, 11010-1-AP), anti-GABARAP-L1 (WB, Abcam, ab86497), anti-vinculin (Sigma, V9131-100UL), anti-ATG2A (Proteinech, 23226-1-AP) and ATG2B (Proteintech, 25155-1-AP), LAMP-2 (DSHB, clone H4B4), c-Myc (DSHB, clone 9E10).

### Cell culture and reagents

HEK293T and U2OS or U2OS ATG2A/B double knock-out cells were maintained in Dulbecco’s modified Eagle’s medium (DMEM; Invitrogen 10313021) supplemented with 10% foetal bovine serum (FBS), 5 U/ml penicillin and 50 μg/ml streptomycin, 1mM L-glutamine and 1% sodium pyruvate. For starvation in nutrient□deplete medium, the cells were incubated 2 h in Earle’s balanced salt solution (EBSS; Gibco, 24010□043). Bafilomycin A1 (BafA1; Enzo, BML□CM110□0100) was used at 200 nM. ATG2A/2B DKO cells were stably transfected by retroviral transduction of pMSCV-Flag-HA (iTAP) vectors or lentiviral transduction of pHAGE-NTAP-mCherry or pHAGE-N-EGFP vectors. Briefly, HEK293T cells were transfected with iTAP vectors with pCG-GagPol and pCG-VSVG (retrovirus) or pHAGE constructs in combination with pCMV□VSV□G and psPAX2 (lentivirus) packaging vectors. Virus containing media was harvested 48 h post□transfection, centrifuged at 300xg, passed through a 0.450□μm filter and added to ATG2A/B DKO cells in the presence of 10 μg/ml polybrene (Sigma, H9268-5G). Transduced cells were selected by addition of 1 μg/ml Puromycin (iTAP and pHAGE-nTAP-mCherry) or 10µg/ml Blasticidin (pHAGE-EGFP) 48 h after addition of viral media.

### CRISPR/Cas-9 gene editing

CRISPR/Cas9-mediated deletion of ATG2A (NM_015104.3) and ATG2B (NM_018036.7) in osteosarcoma cells (U2OS) was performed by using the Cas9 D10A ‘nickase’ mutant and paired gRNAs approach [53] was used to target exon 1 of both ATG2A (5′-CCATGGTCAAACTGTGTGAAAGA-3′ and 5′-TACTTGCTGCACCACTACTTAGG-3′) and ATG2B (5′-CCGTTTTCGGAGTCCATCAAGAA-3′ and 5′-CCTGCCGGTACCTCCTGCAGAGG-3′). ATG2A and ATG2B-targeting gRNAs were transfected into 1×10^6^ U2OS cells followed by selection with 1 μg/ml puromycin for 48hrs, re-transfection, recovery (in puromycin free media) and single cell sorting to isolate clone candidates with the gene deletion.

Endogenous GFP-tagged ATG2A knock-ins were generated using a modified ‘nickase’ strategy (as above). Optimal sgRNA pairs were identified and chosen on the basis of being as close as possible to the point of GFP insertion while having a low combined off-targeting score (ATG2A-sgRNA1: 5’-GTCAAACTGTGTGAAAGAGC-3’ & sgRNA2: 5’-AGATGTCACGATGGCTGTGGC-3’). Complementary oligos with BbsI compatible overhangs were designed for each and these dsDNA guide inserts ligated into BbsI-digested target vectors; the antisense guide (sgRNA2) was cloned onto the spCas9 D10A expressing pX335 vector (Addgene plasmid no. 42335) and the sense guides (sgRNA1) into the puromycin-selectable pBABED P U6 plasmid (Dundee-modified version of pBABE-puro plasmid). A donor construct consisting of GFP flanked by approximately 500 bp homology arms were synthesized by GeneArt (Life Technologies); each donor was engineered to contain sufficient silent mutations to prevent recognition and cleavage by Cas9 nuclease. Both sgRNA and donor constructs were transfected into U2OS cells, selected in 1μg/ml puromycin for 48hrs, re-transfected and allowed to recover in puromycin free complete media. When confluent, cells were single cell sorted for GFP-positive populations and homozygous clones selected for further analysis.

### Western blot and immunoprecipitation

Cells (HEK293T, U2OS) were lysed in NP-40 lysis buffer (50mM TRIS, pH7.5, 120mM NaCl, 1% NP-40) supplemented with Complete® protease inhibitor (Roche) and phosphatase inhibitor cocktail (Roche). Lysates were passed through a 27G needle, centrifuged at 21000xg and incubated with either anti-GFP agarose (Chromotek, gta-20) or anti-HA agarose (Sigma A2095) washed 3 times in lysis buffer and subjected to SDS-PAGE and western blot. For total cell lysis (TCL), cells were lysed in 50mM Tris pH 7.5, 150mM NaCl, 1mM MgCl_2_, 1% SDS. TCL buffer was supplemented with Complete® protease inhibitor (Roche), phosphatase inhibitor cocktail (Roche) and Benzonase (VWR/Fischer scientific) at 1µl per ml buffer. Samples were boiled in 3x Laemelli buffer prior to SDS-PAGE. Unless otherwise stated, NuPAGE™ 4-12% Bis-Tris gradient gels (Invitrogen) were used. Gels were transferred onto activated PVDF membranes (Immobilon Psq, 0.2µm, Merck) prior to blocking and incubation with the indicated primary antibodies.

### Autophagy flux by flow cytometry assay

U2OS ATG2A/2B DKO cells were transfected with mCherry-EGFP-LC3B tandem tagged reporter construct [45, 54], grown in G418/Neomycin selection (800µg/ml) and single cell cloned. U2OS-ATG2A/B DKO-Tandem LC3B cells were then transduced with retrovirus containing iTAP ATG2A constructs and selected for in G418 + 1.5ug/ml puromycin and stable cells generated. These were treated as indicated, scraped in PBS and fixed in 4% PFA for 15mins, washed and then subjected to flow cytometry analysis. All flow cytometry experiments were at least three times using 10,000 cells per cell line per treatment. The cells were then analysed and sorted on an LSR Fortessa (Becton Dickinson) flow cytometer. cells were gated according to forward scatter and side scatter and dead cells were excluded from analysis. GFP fluorescence measured by excitation at 488 nm and emission detected at 530 ± 30 nm and mCherry fluorescence measured by excitation at 561 nm and emission detected at 610 ± 20 nm. Flow data was analysed using FlowJo software.

### Immunofluorescence & confocal microscopy

Cells grown 18mm glass coverslips, were treated as described and subsequently fixed in 4% paraformaldehyde/PBS (PFA; Santa Cruz, 30525-89-4) for 10 min at room temperature and washed 3x in PBS. Cells were then washed in PBS/0.1% saponin twice and primary antibodies incubated for 1 hr at room temperature in 5% BSA/PBS/0.1% saponin. DAPI (Molecular Probes) was added during primary antibody incubation. Coverslips were then washed twice in PBS/0.1% saponin and secondary antibodies (Invitrogen Donkey Anti-Mouse, -Rabbit, -Rat Alexa dyes (488, 555, 647) were used in combination depending on the primary antibody species and incubated in PBS/5% BSA/0.1% saponin. For detection of endogenous GFP-ATG2A, nanobody boosters towards GFP (anti-GFP, Atto-488 coupled, Chromotek; gba488-100) were used to enhance the signal. Secondary antibodies were then washed twice in PBS/0.1% saponin, once in PBS and once in ddH_2_O to remove the residual saponin prior to mounting in ProLong Diamond Antifade containing Mowiol (Invitrogen, p36965). Cells were imaged using a Zeiss 710 confocal microscope with a 63× objective lens. Subsequent image analysis was performed using FIJI (ImageJ) [55].

### Lipid droplet induction and imaging

Cells were set up on glass coverslips and incubated with either complete medium plus 2% BSA or complete media plus 2% BSA/500µM oleic aid for 16 h. Cells were then fixed in 4% PFA/PBS for 10 minutes and permeabilised using the saponin method detailed above. Coverslips were incubated with anti-HA primary antibody and donkey anti-rat Alexa 657 secondary antibody. Lipid droplets were stained using 5µM BODIPY 493/503 (Thermo Fisher Scientific) to stain neutral lipids. Samples were mounted, imaged and analysed as detailed above.

### Protein expression and purification

GST-tagged mammalian ATG8 fusion proteins were cloned into pGEX-4T-1 (GE Healthcare) and expressed in *Escherichia coli* BL21 (DE3) cells in LB medium as previously described [5]. Expression was induced by addition of 0.5 mM IPTG and cells were incubated at 16°C overnight. Harvested cells were lysed using sonication in a lysis buffer (20 mM Tris-HCl pH 7.5, 10 mM EDTA, 5 mM EGTA, 150mM NaCl) and the supernatant was subsequently applied to Glutathione Sepharose 4B beads (GE Healthcare). After several washes, fusion protein-bound beads were used directly in GST pulldown assays.

### Proteinase K protection assay

Proteinase K assay was performed as previously detailed[54]. Briefly, U2OS ATG2A/B DKO or DKO plus ATG2A-WT, ATG2A-mLIR or ATG2A-mYFS cells were grown in complete media or starved (EBSS) in the presence of BafilomycinA1 (200nM) for 4 h, scraped in PBS and centrifuged at 500 x g. The cells were then resuspended in PBS/6.5μg/ml digitonin incubated for 5 min at room temperature and then for a further 30 min on ice. Samples were subsequently centrifuged at 13,000 x g and the supernatant removed. The membrane fractions were then resuspended in 50mM Tris, pH 7.5, 0.18M Sucrose. Resuspended membrane pellets were then incubated with either buffer only, buffer + 100ng/ml proteinase K (PK) or PK+ 0.1% Triton X-100 (PK+TX) for 10min at 30°C. The reaction was stopped by addition of 3x Laemmli sample buffer and boiled at 95°C.

### Cloning and plasmid generation

pDONOR-ATG2A and pDONOR-ATG2B were kind gifts from C.Behrends. These were used in conjunction with Gateway cloning system (Invitrogen) pDEST-CMV-Myc and pMSCV-Flag-HA-IRES-Puro (iTAP) to generate plasmids expressing either ATG2A or ATG2B. Site-directed mutagenesis was carried out to mutate the wild-type gene for the required amino acid substitutions.

### Plasmids

| Plasmid/epitope tag | Gene/mutation | Reference |
| --- | --- | --- |
| pDONOR223-hATG2A | Human ATG2A | This study |
| pDONOR223-hATG2B | Human ATG2B | This study |
| (iTAP) pMSCV-Flag-HA-ATG2A | ATG2A-WT | This study |
| (iTAP) pMSCV-Flag-HA-ATG2A-mLIR | ATG2A aa1362-1365 FCIL/AAAA | This study |
| (iTAP) pMSCV-Flag-HA-ATG2A-mYFS | ATG2A aa1395-1397 YFS/AAA | This study |
| pDEST-CMV-Myc-ATG2A-WT | ATG2A-WT | This study |
| pDEST-CMV-Myc-ATG2A-mLIR#1 | ATG2A aa536-539 YTEI/AAAA | This study |
| pDEST-CMV-Myc-ATG2A-mLIR#2 | ATG2A aa926-929 FSTL/AAAA | This study |
| pDEST-CMV-Myc-ATG2A-mLIR#3 | ATG2A aa981-984 YFCL/AAAA | This study |
| pDEST-CMV-Myc-ATG2A-mLIR#4 | ATG2A aa1092-1095FLDV/AAAA | This study |
| pDEST-CMV-Myc-ATG2A-mLIR #5 | ATG2A aa1362-1365 FCIL/AAAA | This study |
| pDEST-CMV-Myc-ATG2B-WT | ATG2B-WT | This study |
| pDEST-CMV-Myc-ATG2B-mLIR#1 | ATG2B aa845-848 WPRI/AAAA | This study |
| pDEST-CMV-Myc-ATG2B-mLIR#3 | ATG2B aa979-982 FENI/AAAA | This study |
| pDEST-CMV-Myc-ATG2B-mLIR#4 | ATG2B aa1026-1029<br>FSTV/AAAA | This study |
| pDEST-CMV-Myc-ATG2B-mLIR#5 | ATG2B aa1306-1309<br>YVRV/AAAA | This study |
| pDEST-CMV-Myc-ATG2B-mLIR#6 | ATG2B aa1491-1494 FCIL/AAAA | This study |
| pGEX-4T1 alone | GST only | [5] |
| pGEX-4T1-LC3A-ΔG | MAP1LC3A deletion of c-term Gly | [5] |
| pGEX-4T1-LC3B-ΔG | MAP1LC3B deletion of c-term Gly | [5] |
| pGEX-4T1-LC3C-ΔG | MAP1LC3C deletion of c-term Gly | [5] |
| pGEX-4T1-GABARAP-ΔG | GABARAP deletion of c-term Gly | [5] |
| pGEX-4T1-GABARAP-L1-ΔG | GABARAP-L1 deletion of c-term<br>Gly | [5] |
| pGEX-4T1-GABARAP L2-ΔG | GABARAP-L2 deletion of c-term<br>Gly | [5] |
| pEGFP-C1 empty | GFP only | [5] |
| pEGFP-C1-LC3B | GFP-tagged MAP1LC3B | [5] |
| pEGFP-C1-GABARAP | GFP-tagged GABARAP | [5] |
| pMRXIP-GFP-Stx17 WT | GFP tagged Syntaxin 17 | [48] |

## Abbreviations

ATG2: Autophagy-related protein 2
ESCRT: endosomal sorting complex required for transport
GABARAP: Gamma-aminobutyric acid receptor-associated protein
GIM: GABARAP Interaction Motif
GST: Glutathione S-transferase
LIR: LC3 interaction region
MAP1LC3 (LC3): Microtubule-associated proteins 1A/1B light chain
PtIns3P: phosphatidylinositol-3-phosphate
PI3K: PtsIns3P kinase
SQSTM1: Sequestosome-1
VPS34: Vacuolar Protein sorting 34
WIPI: WD repeat domain phosphoinositide-interacting protein

## Acknowledgements

We would like to acknowledge P.Crocker, I. Ganley, A.Gubas, I.Dikic and K.Ryan for critical reading of the manuscript and valuable insights and discussions. We are grateful to Prof. Sharon Tooze for providing a kind gift of WIPI2 and WIPI4 antibodies as well as WIPI expression plasmids. The anti-LAMP-2 (H4B4) and anti-c-Myc (9E10) antibodies were obtained from the Developmental Studies Hybridoma Bank, created by the NICHD of the NIH and maintained at The University of Iowa. We gratefully acknowledge the support from ERASMUS+ for the support of MB and LvdB. This work was supported by a grant from Tenovus Scotland (T16/44).

## Author contributions

MB, LVDB, NG and ND performed experiments, TJM designed and synthesised CRISPR/Cas9 guides, ARP assisted in confocal imaging and DGM designed, performed and analysed experiments. DGM wrote the manuscript.

## Conflict of interest

The authors declare that they have no conflict of interest.

## Expanded View

**Figure EV1.**
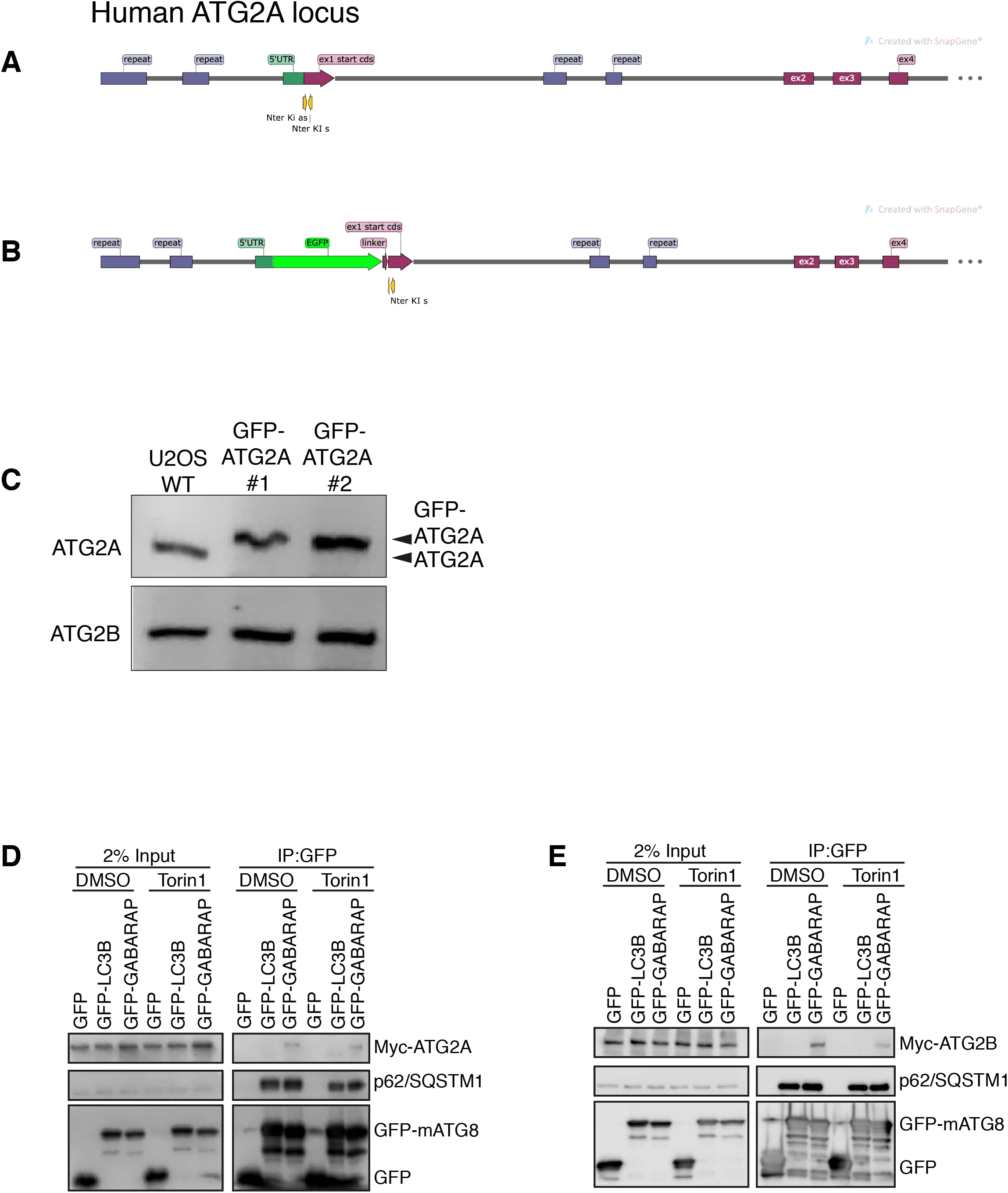
Generation of GFP-tagged endogenous ATG2A. **(A)** Strategy for insertion of GFP-tag upstream of human ATG2A exon 1. Graphic shows position of guides and locus before and after **(B)** GFP-tag plus linker insertion. **(C)** Western blot of total cell lysates from parental wild type (WT) and GFP-ATG2A CRISPR/Cas9 Knock-in clones 1 and 2 using anti-ATG2A and Atnti-ATG2B antibodies. **(D)** Myc-tagged ATG2A-wild type (WT) or ATG2B-WT **(E)** was co-expressed with GFP-alone, GFP-LC3B or GFP-GABARAP in HEK293T cells, lysed and anti-GFP nanobodies used to immunoprecipitate GFP-tagged proteins. Samples were subjected to western blotting and probed for the presence of Myc-ATG2A in immunoprecipitated samples. Anti-p62/SQSTM1 was used as an internal control for the immunoprecipitated samples.

**Figure EV2.**
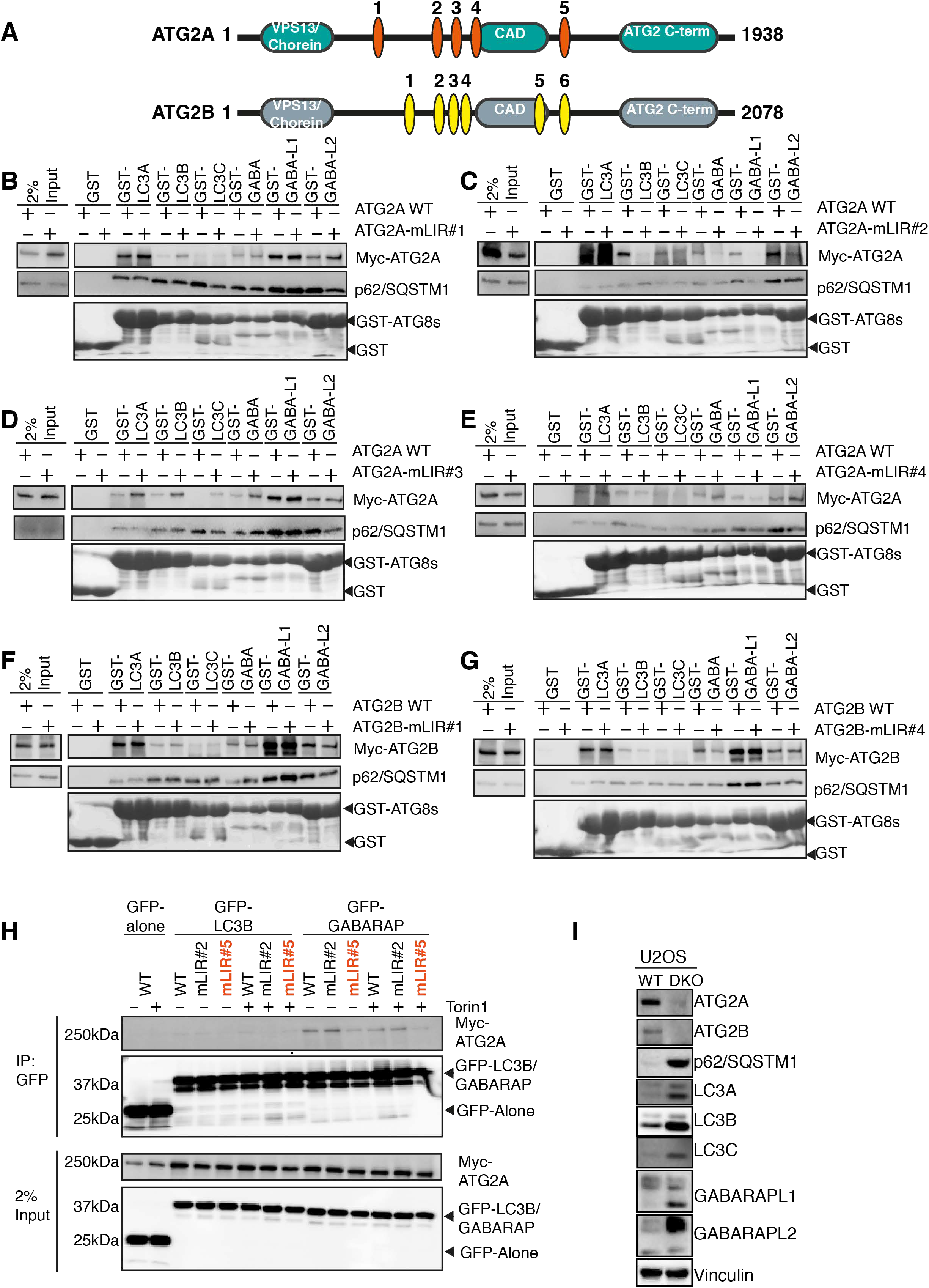
ATG2A and ATG2B contain a highly conserved LC3 Interaction Region (LIR). **(A)** Domain structure of ATG2A (green) and ATG2B (grey) proteins. Both ATG2s contain an N-terminal VPS13/Chorein domain, ATG2 C-terminal autophagy domain (CAD motif) and ATG2 c-terminal domain. Position and sequence of putative ATG2 LC3 interaction regions (LIRs) as identified by iLIR and manual annotation. ATG2A has 5 potential LIRs and ATG2B has 6 potential LIRs. See Table 1 for details. **(B)** Myc-tagged ATG2A-wild type (WT) and putative LIR mutants, where the potential core motif was mutated to alanine were used in a pull-down assay with GST-tagged mammalian ATG8 proteins. Shown are ATG2-mLIR#1, ATG2-mLIR#2 **(C)**, ATG2-mLIR#3 **(D)** and ATG2-mLIR#4 **(E)**. ATG2-mLIR#5 is shown on Figure 2A. ATG2A-WT or -mLIRs were over expressed in HEK293T cells, and lysates were incubated with purified GST alone or GST-tagged LC3A, -LC3B, LC3C, -GABARAP, -GABARAP-L1 or GABARAP-L2. Samples were spun, washed and blotted for the presence/absence of Myc tagged ATG2A using anti-Myc antibody. Anti-p62/SQSTM1 was used as an internal control for the GST-pull-down samples GST-proteins were visualised by Ponceau S staining of membranes. **(F)** As in (B) but using Myc-tagged ATG2B-WT or ATG2B-mLIR proteins. Shown are ATG2B mLIR#1 and ATG2B mLIR#4 (**G**). ATG2B-mLIR#2 was present on an alpha helix, whereas ATG2B-mLIR#3 and ATG2B-mLIR#5 were not expressed. ATG2B-mLIR#6 is shown on Figure 2A. See also Table 1. Anti-p62/SQSTM1 was used as an internal control for the GST-pull-down samples. All blots are representative of at least n=3 independent experiments

**Figure EV3.**
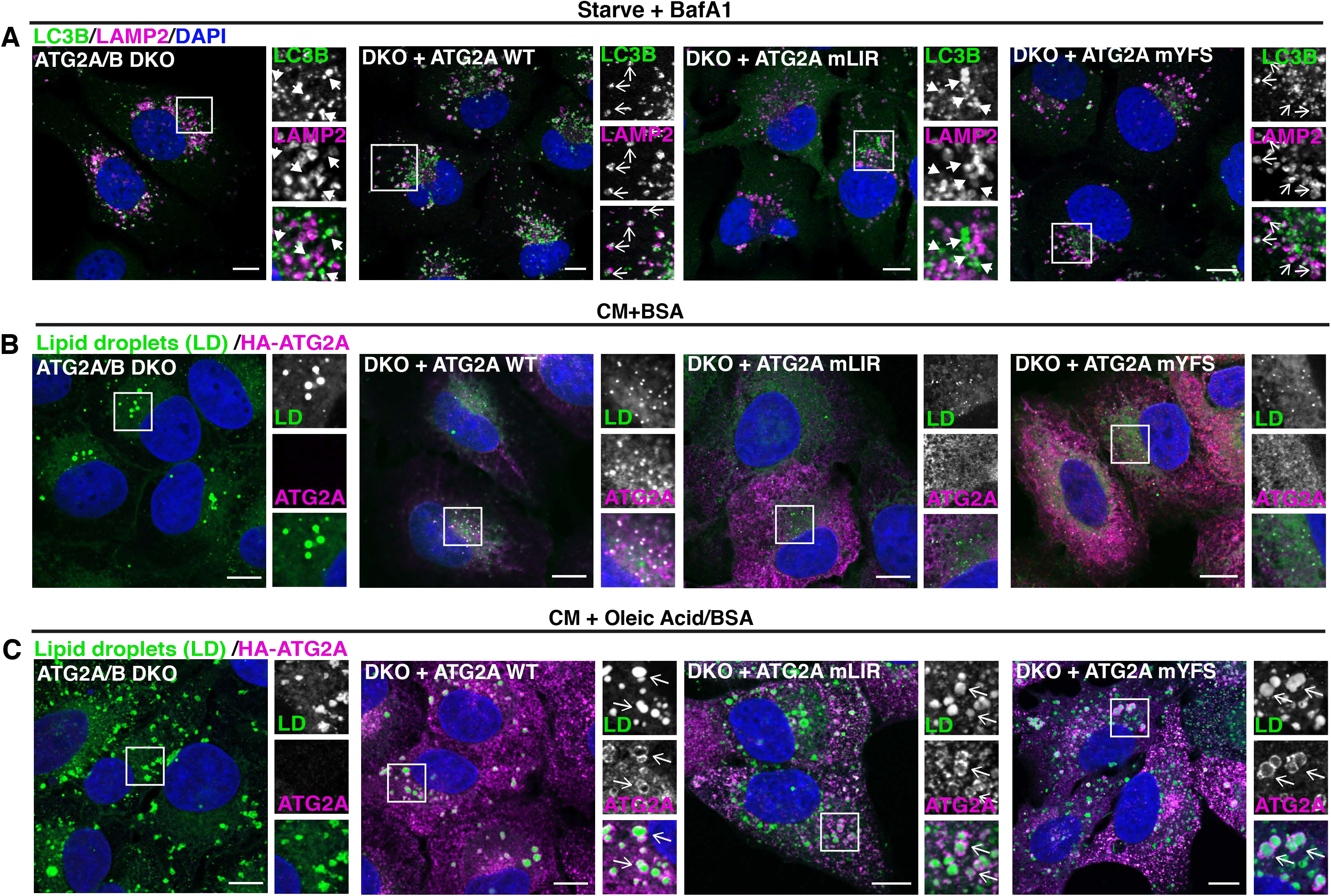
ATG2A-WT and mutants effectively localize to lipid droplets. **(A)** U2OS ATG2A/B DKO cells reconstituted with vector only, HA-ATG2A-WT, -mLIR and - mYFS were stimulated starvation (EBSS) plus BafilomycinA1 (BafA1, 200nM) for 4 hours to stimulate autophagosome generation and prevent their degradation in the lysosome. Cells were fixed and immune-stained for LC3B (magenta) and LAMP2 to visualise lysosomes. DAPI was included (blue) to mark the DNA/nucleus. Closed arrows (ATG2A/B DKO and DKO + ATG2A mLIR) highlight aggregate structures. Open arrows (DKO + ATG2A-WT and ATG2A-mYFS) highlight LAMP2/LC3B positive vesicles. Scale bar 10µm. **(B)** U2OS ATG2A/B DKO cells reconstituted with vector only, HA-ATG2A-WT, -mLIR and - mYFS were stimulated with either 2% BSA only or **(C)** 2%BSA plus 500µM oleic acid for 16 h prior to fixation in 4% PFA. Cells were permeabilised using saponin and stained with anti-HA (ATG2A; magenta) and 5µM BODIPY 493/503 to visualise lipid droplets (green; LDs). DAPI was included (blue) to mark the DNA/nucleus. Arrows arrows (ATG2A/B DKO and DKO + ATG2A mLIR) highlight aggregate structures. Open arrows highlight ATG2A positive lipid droplets. All images are representative of at least n=3 independent experiments. Scale bar 10µm.

**Figure EV4.**
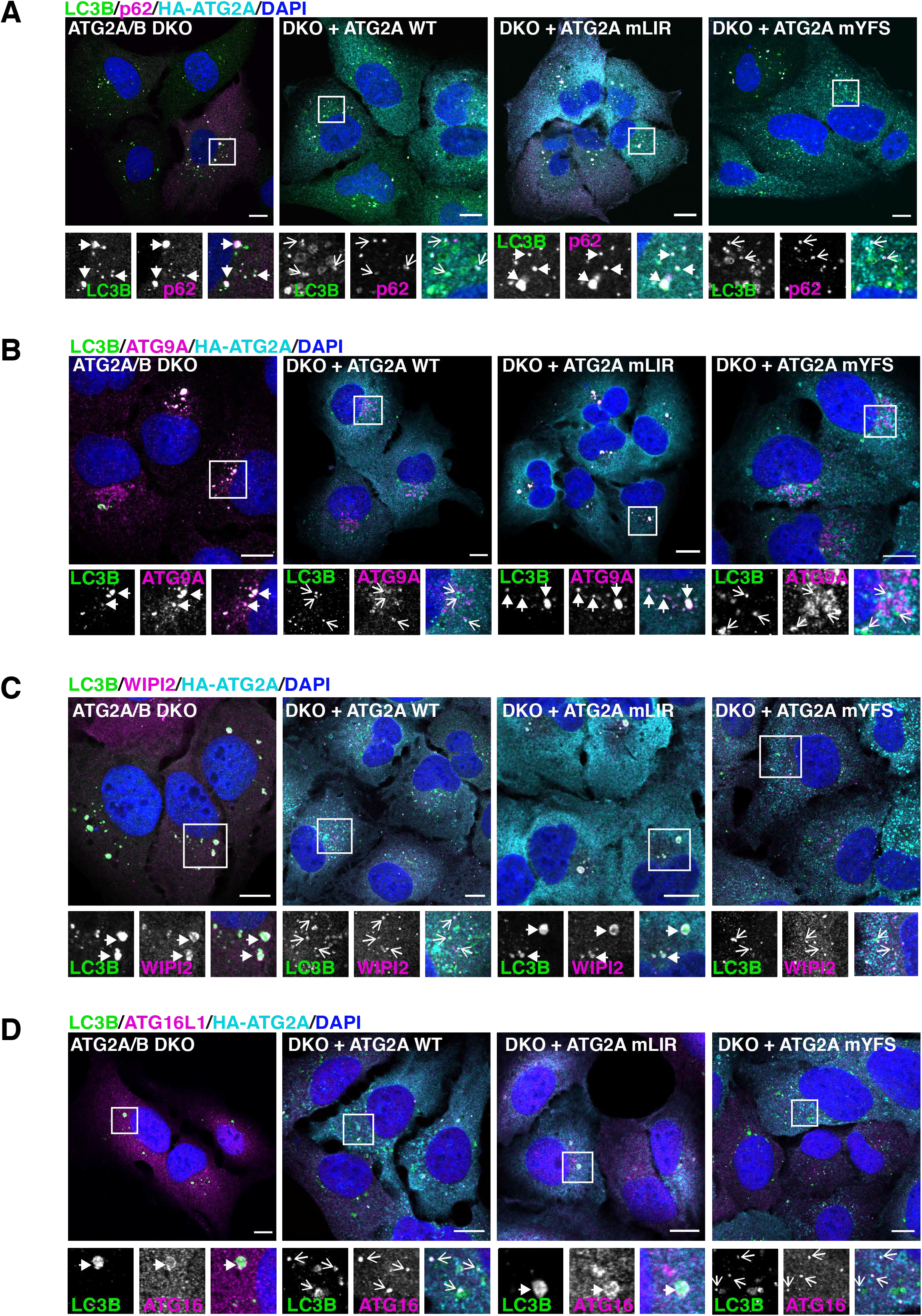
ATG2A-mLIR causes accumulation of early autophagy markers. **(A)** U2OS ATG2A/B DKO cells reconstituted with vector only, HA-ATG2A-WT, -mLIR and - mYFS were stimulated starvation (EBSS) 2 hours to stimulate autophagosome generation. Cells were then fixed and immune-stained for LC3B (green), p62/SQSTM1 (magenta) or **(B)** ATG9A, **(C)** WIPI2 and **(D)** ATG16L1 (Magenta). Anti-HA (ATG2A expression) (cyan) and DAPI (blue; DNA/nucleus) were included. Closed arrows (ATG2A/B DKO and DKO + ATG2A mLIR) highlight LC3B positive aggregate structures. Open arrows (DKO + ATG2A-WT and ATG2A-mYFS) highlight LC3B positive vesicles. All images are representative of at least n=3 independent experiments. Scale bar 10µm.

